# Modelling termites’ tunnelling and decision-making behaviors

**DOI:** 10.1101/2023.10.05.561127

**Authors:** Sajib Mandal, Sebastian Oberst, Joseph C. S. Lai

## Abstract

Termites’ digging and construction behavior plays an important role in understanding termites’ foraging and nesting strategies. Termites build galleries when they tunnel that building behavior is different from ants. Therefore, termite tunnelling behavior such as tunnelling networks (tunnel growth and branching), and direction-changing when generating new branches and encountering obstacles require more research. Measurement of termites’ tunnel growth in the experimental arena is often conducted manually by comparing photo sequences. Here, we observe the tunnelling behaviors of the small-sized desert subterranean termite (*Heterotermes aureus*) and the larger arid land subterranean termite (*Reticulitermes tibialis*) and develop a mathematical model to describe termites’ tunnelling behavior in the arena. The model can estimate the possible tunnel length with respect to termite body length over a certain time based on three data inputs. Another advantage of this model is that it takes only a few seconds to show results. The results of the model are verified numerically. A behavioral model based on a decision tree is also developed to investigate termites’ direction-changing mechanisms in tunnelling when generating branches and encountering obstacles. Thus, this study demonstrates methods to characterize and model termites’ tunnelling mechanism and direction-changing behavior, which might be applicable to other insects’ behavior and algorithm mimicking digging and tunneling behavior of ants.

## I Introduction

Termites are superorganisms and exhibit collective behavior in building nests, tunnelling, foraging food sources, and defending their colony in a group against intruders [1], [2]. For the survival of the colony, tunnelling strategies play a very important role in building their nests/mounds, avoiding enemies, and foraging foods as well [3], [4]. For instance, the tunnel and ventilation shafts of termite nests regulate temperature and gas (oxygen, CO_2_, and methane) levels [5], [6]. Subterranean termites unlock foraging resources by constructing tunnels and galleries, and releasing a trail-following pheromones to provide nestmates with information to track the path from the nest to food source [7]–[9]. The emergence of tunnels and galleries is important to understand the collective behaviors of termites and other eusocial insects [9]–[11].

Tunnelling and branching patterns can vary among species, reflecting species-specific digging and construction behaviors [12], [13]. Even groups of termites of a certain species can exhibit differently emerging patterns in experimental arenas and individual behavior may not correspond directly to that of the group [14]. In tunnelling, the excavator digs sand with its mandibles, forms it into a ball with legs, passes it backward to the individual behind it, and moves forward to expand the tunnel [14]. Termite deposition behavior in tunnelling varies over tunnels’ widths and the deposited particles’ functions [15]^1^. Branching in tunnels is representative of the termites’ search for food sources [16]. Most studies have been focused on the development of tunnel and branch patterns, but little is known about how they make decision in the tunnelling and branching and how they change tunnel and branch direction when they face obstacles. To describe termites’ behavior when they tunnel a decision tree is used.

To observe termites’ tunnelling behaviors in lab experiments, researchers mostly set up two-dimensional experimental arenas with transparent layers and capture photos at regular time intervals or continuous videos for further analysis. Some previous studies investigated the overall termites’ activities in the experimental arena by using program languages such as Python and C/C++ [17]–[19]. These systems automatically load photos and analyze them, but require standardized setups used with high-quality photos having an identical number of pixels, and the interpretation of results is sometimes not accurate. Most studies have directly measured termite tunnel and gallery length [13], [14], and shelter tube (gallery) construction dynamics [20] by categorizing the whole system into small groups on geometric patterns. Very few studies are concerned with the structure itself or its connectivity [21], [22]. Since the identification of the groups and measurement of the systems is performed arbitrarily by researchers, results are intuitive and can vary. Recently, Mizumoto (2023) developed a software using Python to measure termite tunnel length manually based on photo analysis [23]. It estimates the tunnel length by detecting the change in subsequent photos. The disadvantages of this system are that it requires preparation of standardized photos (i.e., the same position and same pixels) which is time-consuming, and sometimes cannot detect small changes in the tunnel growth due to the low resolution of the camera. Therefore, there is a need to develop a model that can help the manual measuring process, reduce the effort and time, and predict the next possible outcome. In this case, the finite difference method might be helpful in developing such a model to predict the results based on the previous data [24], [25].

We develop a mathematical model to investigate termites tunnelling behavior in experimental arenas inspired and extracted by Mizumoto (2023) [23]. This model can interpret how tunnel length grows and estimates the possible tunnel length over time. Besides, the model generates results based on only three data inputs in a few seconds and can predict the next possible outcomes. Since it requires only three data, researchers need to estimate the total tunnel length only three times so users need not standardize all photos, thus they can save time. To describe termites’ tunnelling and decision-making behavior in the experimental arena, we develop a behavioral model based on a decision tree. This model describes the generation of branches and the selection of tunnel directions by termites when they meet the boundary of the experimental arena. The numerical results of the proposed models are compared with experimental studies. This study represents the application of mathematics in insect biology. The limitations and future directions of this study are finally discussed.

## II. Materials and Methods

### A. Experimental Design and Data

To develop a mathematical model and a decision tree to describe termites tunnelling over time, we collected experimental data provided by Mizumoto et al. (2020) [14] and Mizumoto (2023) [23]. We observed the tunnelling of *Heterotermes aureus* and *Reticulitermes tibialis. H. aureus* colonies were collected from cholla and mesquite deserts in Gila and Maricopa Countries, Arizona [14] and *R. tibialis* colonies were collected from a pine forest in Pinal Country, Arizona [14]. The experimental arena had a diameter of about 125 mm, and had three layers: the bottom layer was made with acrylic board, the middle layer was filled with moist white sand (10% distilled water by volume) with 1mm thickness based on termite head size, and the top layer was a transparent Perspex sheet [14]. A teardrop-shaped entrance was considered for termites to enter the arena and a glass plate was used to cover the area as presented in Fig.1A and 20 worker termites were used for each experiment [14]. The average body size of *H. aureus* and *R. tibialis* were 3.9 mm and 4.4 mm, respectively [14]. Therefore, the arena was 32 times of *H. aureus* body size and 28 times of *R. tibialis* body size [14]. We considered 15 experimental results (three colonies × five replicates) for both *H. aureus* and *R. tibialis* [14]. Figs. 1B and 1C display respectively a sample of the tunnelling behavior of *H. aureus* and *R. tibialis* in the arena.

**Fig. 1.**
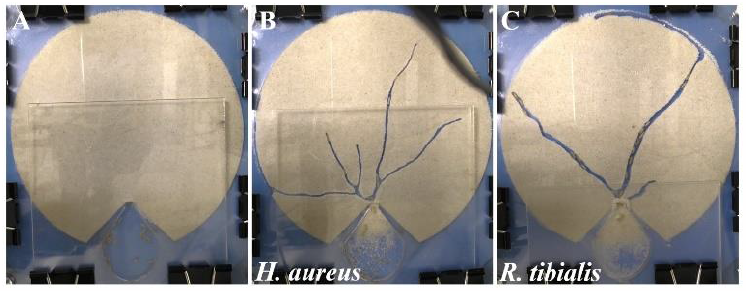
(A) Initial stage of the experimental arena (*t=0h*), (B) tunnelling pattern of *H. aureus* in the arena (H-B1, *t =*20 h), and (C) tunnelling pattern of *R. tibialis* in the arena (R-A5, *t=*20h) [23].

Figs. 2 and 3 show the total tunnel length compared to the body size of *H. aureus* and *R. tibialis*, respectively for 15 experiments and their average tunnel length [23]. Here, the total tunnel length stands for the sum of all tunnel lengths in the arena and the tunnel length was measured in millimeters (mm) for the first 20 hours. Termites tunnelling behaviors vary over species and arenas i.e., termites’ groups presented in Figs. 2 and 3. They also show that some termite groups tunneled up to three times more than some other groups and tunnelling is unpredictable. According to Figs. 2 and 3, tunnels grow significantly between the start and approximately the 5^th^ hour for almost all arenas and then slowly decrease over time. The figures show that *R. tibialis* tunnels 9.9% more (and has wider tunnels) than *H. aureus* in average due to 0.5 mm larger in body size than *H. aureus*.

**Fig. 2.**
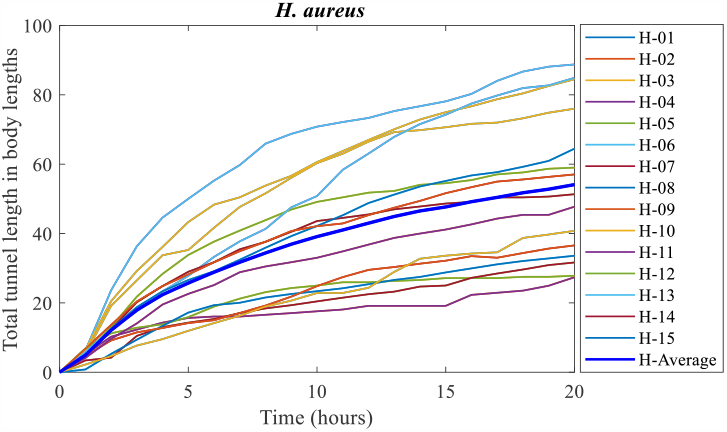
Total tunnel length in body lengths for the termite *H. aureus* estimated from the results of the experiments [23]. Here, B, C, and D represent colony numbers. Statistical data are provided in Supplementary Information 1.

**Fig. 3.**
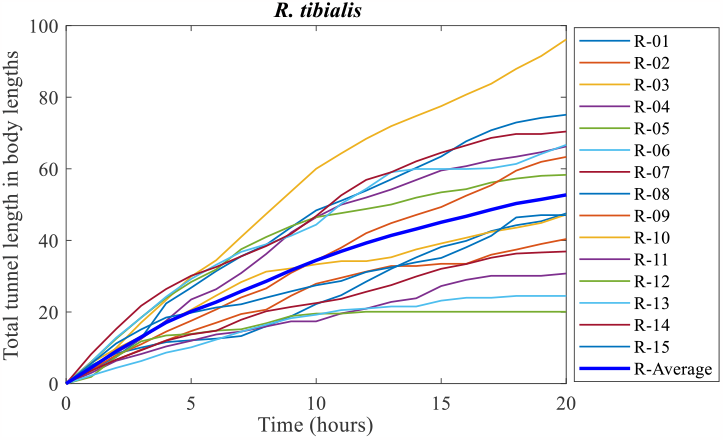
Total tunnel length in body lengths for the termite *R. tibialis* estimated from the experiments results [23]. Here, A, B, and C represent colony numbers. Statistical data are provided in Supplementary Information 2.

### B. Finite Difference Equation

A finite difference equation represents a functional equation involving a time/position varying variable and a finite difference operator. Finite differences are widely used in numerical analysis to approximate derivatives [24], [25]. The solution methods of the difference equation are almost similar to those of differential equations. Here, out of the three types of difference methods (forward, backward and central), the backward difference method is used because it can predict the next outcomes based on the previous data and generate a series of results over time [24], [25]. To define the backward difference equation, let *x* be the variable, *h* be a constant, and Δ_*h*_ represent the difference operator which maps a function *f* to the function Δ_*h*_[*f*][25] . Then the backward difference equation can be defined as

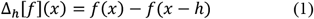

If the difference between two sequential terms is 1, e.g., *x*_1_, *x*_2_, *x*_3_, … … , *x_n_*, *h* can be replaced by 1 and then the Eq. (1) can be written as

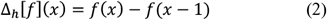

### C. Model Description

Let *x*(*t*) be the time-varying variable that represents the tunnel length over time *t* = 0,1,2,3, … … , *n* . To develop a mathematical model to describe the tunnel growth in termites body lengths, three terms are used as inputs to the system: the first, third and fifth terms i.e., *x*(1), *x*(3) and *x*(5). Since the total tunnel length grows more at the initials and then slightly slows down over time [23], therefore, to find more accurate results for the model, the second term that represents the tunnel length at *t* = 2 is defined by 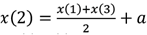, and the fourth term is defined by 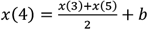, where *a* and *b* are the balancing constants with the value *a* = 2 and 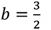. The value of *a* and *b* are determined from the growth of the tunnels observed in the experimental results [23], cf Figs. 2 and 3. In this case, the differences between the average of *x*(1) and *x*(3) and the experimental results of *x*(2) are calculated and then the most commonly fitted value is considered for *a* to get the almost actual value of *x*(2), and the similar process applied to find the value of *b*. Let’s consider the growth of the total tunnel length starting to slow down from the sixth term. The next *n* terms are split into two equations as follows.

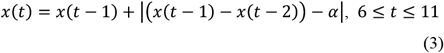

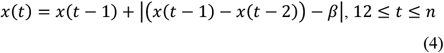

Therefore, Eqs. (3) and (4) describe the next tunnel length as the sum of the previous tunnel length and the absolute tunnel growth per unit time. Here, *α* and *β* are control parameters that balance the growth of tunnel over time with respect to the previous term, and can be defined as follows.

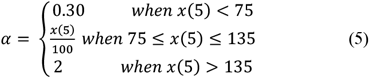

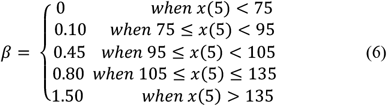

The values of *α* and *β* are determined by observing the changes in total tunnel lengths over time, cf Figs. 2 and 3, and assigning the conditional values based on the value of *x*(5) which predict the results with low errors [23].

Therefore, the tunnel length at any time *t* can be represented as

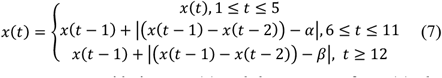

Hence, considering Eq. (7) and the concept of Eq. (1), the growth of total tunnel length in termites’ body lengths can be defined as follows:

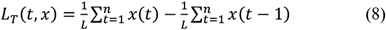

with the initial condition *x*(0) = 0. Here, *L* represents the body length of termite that can vary among species.

Therefore, Eq. (8) is the tunnel length measurement (TLM) model. Eq. (8) can also be defined as 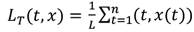. Since, *x*(*t*) represents the tunnel length (not the increment) at time *t*, the second term of Eq. (8) i.e., 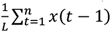 is considered to balance the sum of the tunnel length.

## III. Results of the TLM Model (8)

MATLAB has been used to numerically investigate the findings of the TLM model (8). The main purpose of this section is to examine the findings of the TLM model (8) with experimental results and other previous studies. Running the simulation for the TLM model (8) requires three data inputs: *x*(1), *x*(3) and *x*(5). The time duration and the body size of termite species are specified by the user. The results are expressed in total tunnel length in termites body lengths over time. MATLAB codes are provided in Supplemental Information 3.

Fig. 4 displays the results of the TLM model (8) for time *t* = 20*h* as the same as Figs. 2 and 3. As shown in Fig. 4A, the results of the TLM model (8) compare well with the average tunnel lengths of *H. aureus* and *R. tibialis* determined in the experiments [14], [23]. Fig. 4B shows that the accuracy of the results of the TLM model (8) (in percentage with respect to the average tunnel lengths in body lengths of *H. aureus* and *R. tibialis*, as presented in Figs. 2 and 3) is up to 99.9% for *H. aureus* at *t* = 13 with the lowest being 98.6% at *t* = 2, and 99.8% for *R. tibialis* at *t* = 8, with the lowest being 94.9% at *t* = 13.

**Fig. 4.**
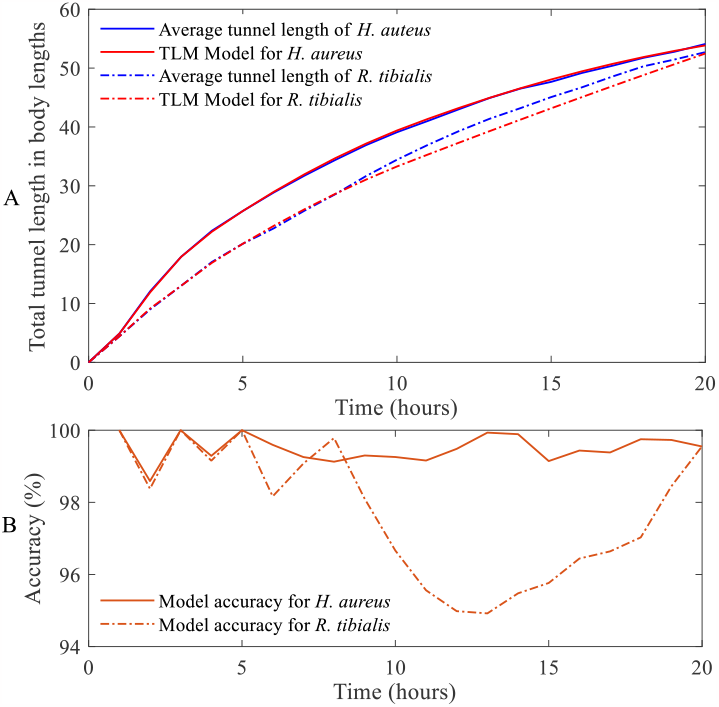
(A) Growth of total tunnel length per body lengths of *H. aureus* and *R. tibialis* over time (hours): comparison between the average tunnel length per body lengths of the termite species and the time series of the TLM model (8), (B) accuracy presented in percentage (%) of the TLM model (8) with respect to the average tunnel length in corresponding body length.

The highest and lowest accuracy of the TLM model (8) are considered for all *x*(*t*), except *x*(1), *x*(3) and *x*(5), as which are inputs taken from the experimental data (hence 100% accuracy) as shown in Fig. 4B. Fig. 4B show that the TLM model (8) can estimate termites tunnel length with up to 99.9% accuracy. Therefore, the results confirm the validity of the TLM model (8).

## IV. Model of Termite Tunnelling

Modelling termites’ tunnelling behavior is difficult because termites’ tunnelling and branching behaviors vary among species and individuals and with environmental factors. Here, we develop a termite behavioral (TB) model to describe termites’ tunnelling behavior in experimental arenas. To develop the TB model, we observed the tunnelling behaviors of *H. aureus* and *R. tibialis* [14], [23] and estimated statistical analysis on their tunnelling and branching behaviors over time (first 2 0*h*). The procedures to develop the model are as follows.

### Procedures Followed to Characterise Termites’ Tunnelling Behaviour and Develop the TB Model

- Names of tunnels and branches: The first four types of tunnels are considered as the Primary, Secondary (Primary branch), tertiary (Secondary branch), and Quaternary (Tertiary branch) tunnels [23]. Tunnel names are assigned by their directions and tunnel/branch numbers. More about naming tunnels are presented in Fig. 5.
- Characterize tunnels and branches: A datasheet provided in Supplementary Information 4) is prepared with tunnel and branch names and statistical analysis is used to characterise tunnelling behaviors.
- TB model development: Calculate the probability of termites’ tunnel selection behaviors for all types of tunnels i.e., primary tunnel to tertiary branch. Two decision trees are developed to describe the termites’ tunnelling and branching behavior, as presented in Figs. 6 and 7.

**Fig. 5.**
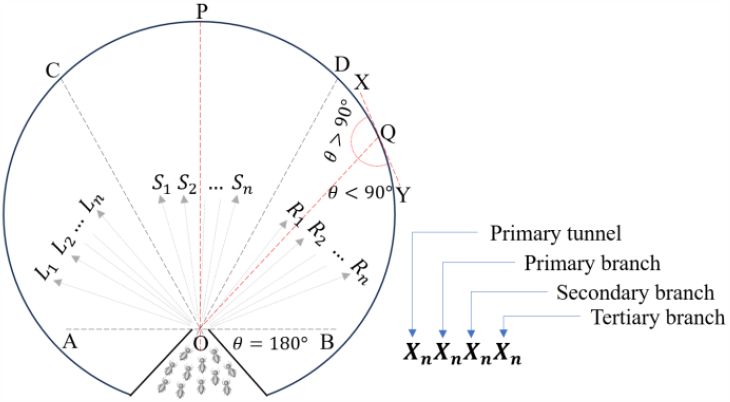
Experimental arena distribution for characterizing tunnels and branches of *H. aureus* and *R. tibialis*. Here, *AOB* represents the horizontal line at the entrance point and the front arena is divided into equal three angles *AOC* = *COD* = *BOD* = 60°. *OQX* and *OQY* are inclined angles at the boundary point *Q* and *XY* is the slope at *Q*. The tunnels within the angles *AOC, COD*, and *BOD* are denoted *L*_1_, *L*_2_, … , *L*_*n*_, *S*_1_, *S*_2_, … , *S*_*n*_, and *R*_1_, *R*_2_, … , *R*_*n*_, respectively. Here, L, S, and R represent Left, Straight, and Right, respectively. These tunnels are called primary tunnels. The tunnels made from the primary tunnels are called primary branches (secondary tunnels), branches from primary branches are secondary branches and similarly for tertiary branches. For example, let’s consider termite starts tunnelling straight i.e., a straight primary tunnel (*S*_1_) and then generates a primary branch at right side (*S*_1_*R*_1_). The primary branch generates a secondary branch at left side (*S*_1_*R*_1_*L*_1_) that generates a tertiary branch at the right (*S*_1_*R*_1_*L*_1_*R*_1_). Therefore, the name of the tertiary branch is *S*_1_*R*_1_*L*_1_*R*_1_ that represent the path from the start.

The behavioral model based on two decision trees describing termites’ decision taking behavior in tunnelling and branching is presented in Figs. 6 and 7. The decision trees are developed based on *H. aureus* and *R. tibialis* tunnelling and branching behavior observed in the experimental arena [14], [23]. Termites start most primary tunnels straight, rather than choosing left or right direction. The possibility of choosing straight direction is more than two times possibility of choosing left or right for *H. aureus* and the possibility of choosing is more than three times for *R. tibialis*, cf Figs. 6 and 7. The straight primary tunnels generate more secondary tunnels/branches and explore more, and it’s true for both termite species. Interestingly, when they start generating branches, they mostly focus on branches than the source tunnels/branches, cf Figs. 6 and 7. The similar behavior of termites is noticed for the left and right primary tunnels. In the case of generating primary branches for both termite species, the possibility to choose left direction is more than other directions. The difference noticed among the three primary tunnels is that the strength primary tunnel generates up to tertiary branches, whereas the left and right primary tunnels stop branching at primary branches for *R. tibialis* and secondary branches for *H. aureus*, which indicates that *H. aureus* build more complex tunnel system by generating more branches compare to *R. tibialis*.

**Fig. 6.**
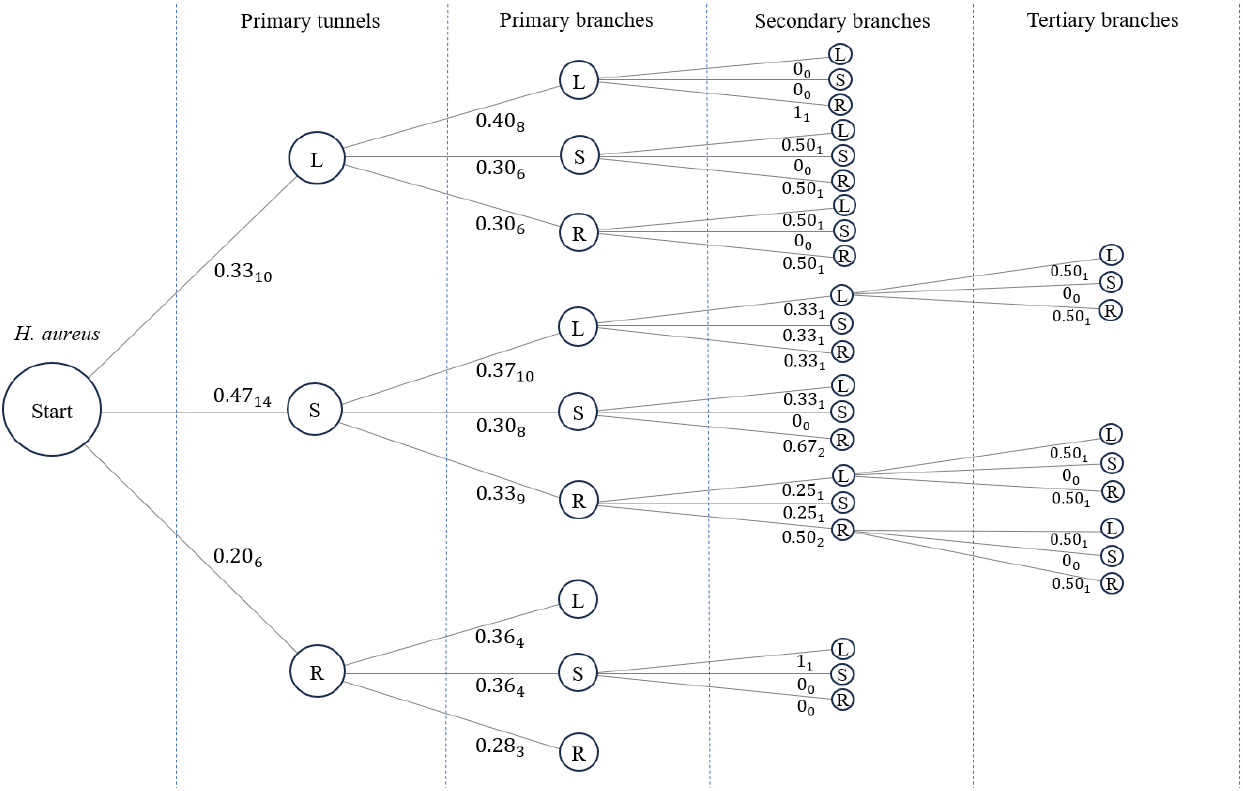
The decision tree describes the tunnelling and branching behaviors of *H. aureus* in experimental arenas. Here, S, L, and R represent Straight, Left, and Right tunnel, respectively. The numeric value shown like “*X*_*n*_” represents the tunnelling intensity, where *X* is the probability (out of 1) of choosing the direction in tunnelling and *n* is the tunnel number. The tunnel number n can vary over the tunnels and branches due to generating more and less tunnels and branches. For instance, *H. aureus* has built n=14 primary straight tunnels (S) and generated 27 (10+8+9) secondary branches. The figure shows *H. aureus* mostly start tunnelling straight and they generate more tunnels/branches in straight and left sides.

**Fig. 7.**
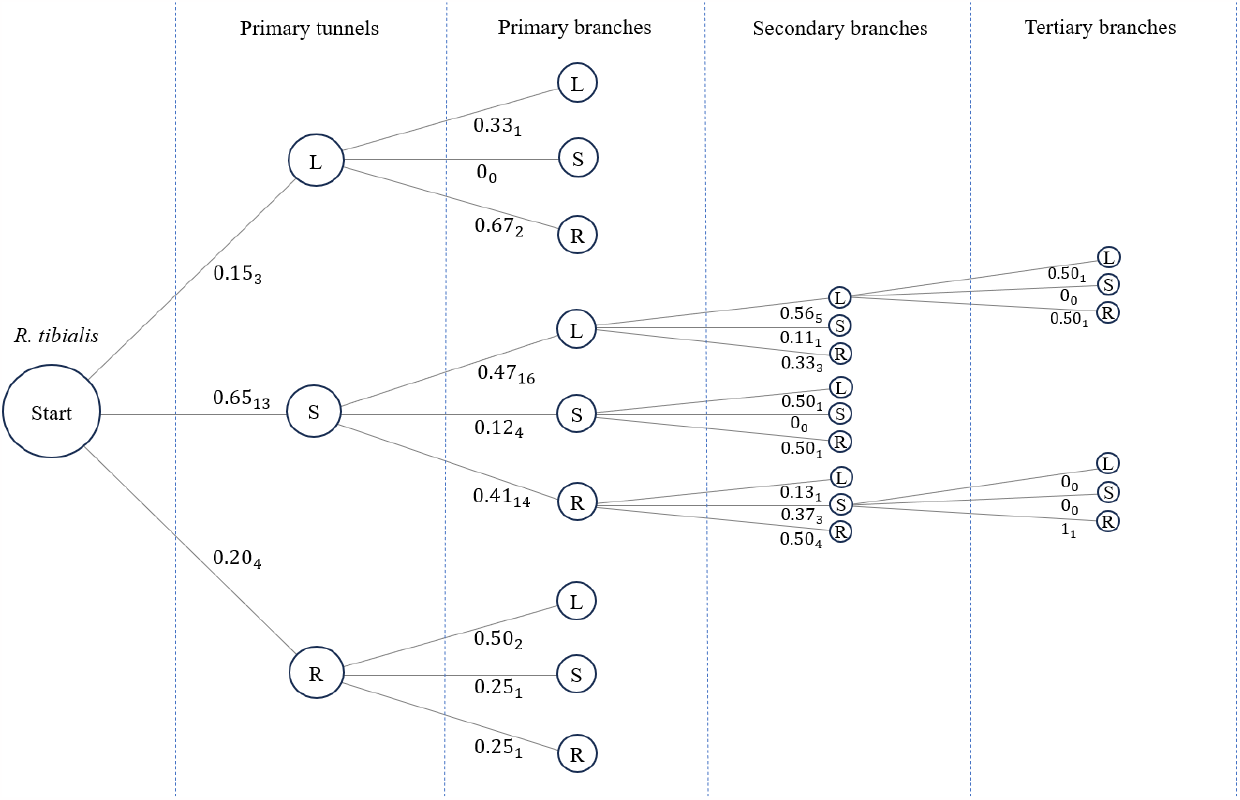
The decision tree describes the tunnelling and branching behaviors of *R. tibialis* in experimental arenas. Here, S, L, and R represent Straight, Left, and Right tunnel, respectively. The numeric value shown like “*X*_*n*_” represents the tunnelling intensity, where *X* is the probability (out of 1) of choosing the direction in tunnelling and *n* is the tunnel number. The tunnel number n can vary over the tunnels and branches due to generating more and less tunnels and branches. For instance, *R. tibialis* has built n=16 primary branches in SL and generated 9 (5+1+3) secondary branches that means 7 SL primary branches have no secondary branches. The tree shows *R. tibialis* has high intensity to start tunnelling straight and the intensity is more than three times of other directions and generates more tunnels/branches, whereas they tunnel and branch less in the left and right sides.

A comparison between the conditional probabilities for *H. aureus* and *R. tibialis* to reach a certain the tunnel and branch are represented in Fig. 8. The figure discloses that H. aureus build more complex tunnel network than R. tibialis by generating more primary and secondary branching, where R. tibialis mostly tunnels straight and generates primary and secondary branched only from straight primary tunnels, cf Fig. 8. The tertiary branches seem almost similar for both the termite species.

**Fig. 8.**
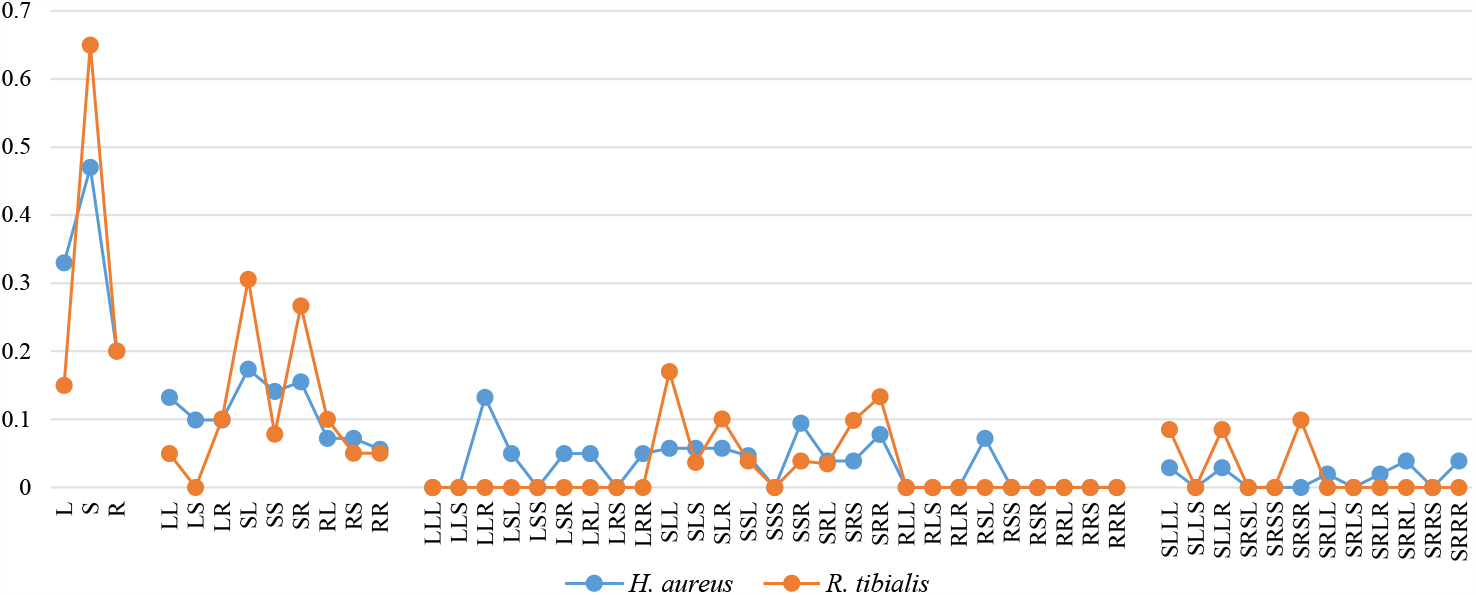
The conditional probabilities for the tunnels and branches of *H. aureus* and *R. tibialis* based on the decision trees, as presented in Figs. 6 and 7. The primary tunnels, primary branch, secondary branch, and tertiary branch are presented horizontally starting with L, LL, LLL, and SLLL, respectively. For tertiary branches, the branches termites explored are considered only. The probability to rely on a certain event is presented vertically which is calculated by multiplying the relative probabilities of the path. The figure shows that *H. aureus* builds a complex tunnel network by generating more primary and secondary branches, whereas *R. tibialis* focus on straight tunnel and builds a tunnel network from the straight primary tunnel(s).

However, two behaviors are observed at the boundary for both termite species: (i) they change direction at either the right or left side, and (ii) they divide the tunnel into two, a right and a left tunnel. They generally chose the Right/Left side where the inclined angle at the boundary is greater than the right angle, i.e., *θ* > 90°, cf Fig. 5. *R. tibialis* always (100%) choose the side where the inclined angle is *θ* > 90°, and *H. aureus* almost (93.3%) follows the same rule. Interestingly, when the inclined angle is *θ*∼90°, they behave randomly and choose either left or right direction, or both. Termites’ behaviors at the boundary are briefly presented in Table 1.

**TABLE I.**
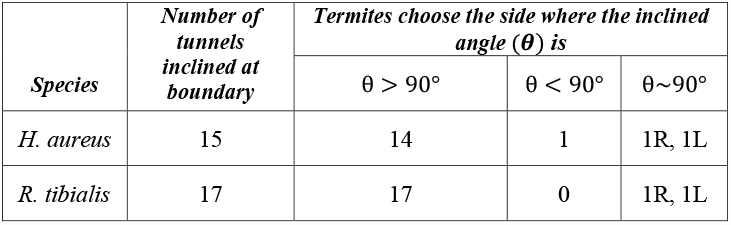
Termites ’ Choice Behaviour at the Arena Boundary Characterised on Inclined Angles [14], [23]. It representes that termites turn the side where the inclined angle is greater than the right angle and can choose left or right when the inclined angle is a right angle or almost a right angle.

| Species | Number of tunnels inclined at boundary | Termites choose the side where the inclined angle ($\theta$) is | | |
| --- | --- | --- | --- | --- |
| | | $\theta > 90^\circ$ | $\theta < 90^\circ$ | $\theta \sim 90^\circ$ |
| *H. aureus* | 15 | 14 | 1 | 1R, 1L |
| *R. tibialis* | 17 | 17 | 0 | 1R, 1L |

## V. Discussion

### TLM Model

A mathematical model is developed to estimate termites’ tunnel length with respect to body length over time. Although this model works on only three data inputs, it can generate almost similar results to the manually estimated tunnel length with high accuracy, as presented in Fig. 4. The total tunnel length needs to be calculated for the three inputs, could be manually or using any tool, thus standardization for all photos such as photo editing, fixing all photos into the same pixels, and naming these sequentially is not required for this model. As a result, researchers can save both time and money to measure termites’ tunnel lengths using this TLM model.

### Behavioral Model

We developed a novel behavioral model based on two decision trees, as presented in Figs. 6 and 7, to describe termites’ tunnelling and branching behaviors i.e., termites’ decision-taking mechanisms in tunnelling and changing directions. Fig. 8 discloses that *H. aureus* builds more complex tunnel network than *R. tibialis* by generating more primary and secondary branches, whereas *R. tibialis* mostly focus on generating and expanding the straight tunnels. One mathematical model is developed to the novel findings is that termites always choose the way at the boundary where the angle is greater than the other i.e., more than a right angle, and randomly any way when the angle is a right angle and almost a right angle, as described in Table 1. Thus, the model provides a clear overview of termites’ possible behaviors in tunnelling and branching in the experimental arena.

### Significance, Limitations, and Future Directions

In this study, we observed the tunnelling and branching behavior of *H. aureus* and *R. tibialis* to develop the TLM model and the TB model. The TLM model generates low errors (0.01% to 5.1%, as presented in Fig. 4B) when it is considered to estimate the average total tunnel length in termites’ body lengths, the error may vary for a single tunnel because a small group of termites generally shows random tunnelling behavior (discrete growth of tunnels) in the experimental arena due to less collective behavior among a handful individuals. Since termites tunnelling behavior changes among species, colonies, groups, and individuals, the accuracy/error of the developed models may vary for other termite species and even other colonies of *H. aureus* and *R. tibialis*. The models can be improved by observing termites tunnelling behavior for the groups of various individual numbers. Termites can behave differently for the variation in arena setup or size. The models also can be further developed by considering a variation of arena setup and/or size.

### Outlooks

The tunnel length measurement (TLM) model is a mathematical model developed for the first time to measure termites’ tunnel length in termites’ body length and the termite behavioral (TB) model is also developed for the first time in this research area to describe termites tunnelling and branching behaviors. This study will assist researchers in developing new models and ideas to contribute to merging mathematical modelling with behavioral insect biology and developing probability models.

## Acknowledgment

This research was supported under the Australian Research Councils (ARC) Discovery Project funding scheme (project No. DP200100358) and the ARC Linkage funding scheme (project No. LP200301196).

## Footnotes

1 Hence ‘tunnelling’ resembles rather a subterranean gallery building.

