## Supplementary Information 1-4 for "Modelling termites’ tunnelling and decision-making behaviors": Supplemental Information 3.pdf

………………………………………………………….………….. TLM Model ……………………………………………………………………..

```
clc
clear all

x(1) = input("Please insert the tunnel length for the 1st hour= ");
x(3) = input("Please insert the tunnel length for the 3rd hour= ");
x(5) = input("Please insert the tunnel length for the 5th hour= ");
n = input("Time= "); %It is better to fix the time to avoid repeating the
input
body = input("Body size of the species= "); %It is better to fix the body
size of the species to avoid repeating the input

x(2) = ((x(1)+x(3))/2)+2;
x(4) = ((x(3)+x(5))/2)+1.5;

if 140<=x(5)
    alpha = 2;
elseif (75<=x(5))&&(x(5)<140)
    alpha = x(5)/100;
else
    alpha = 0.30;
end

for t=4:9
x(t+2) = x(t+1) + abs((x(t+1)-x(t))-alpha);
end

if 140<=x(5)
    beta = 1.5;
elseif (110<=x(5))&&(x(5)<140)
    beta = 0.80;
elseif (95<=x(5))&&(x(5)<110)
    beta = 0.45;
elseif (75<=x(5))&&(x(5)<95)
    beta = 0.10;
else
    beta = 0;
end

for t=10:n-2
    x(t+2) = x(t+1) + abs((x(t+1)-x(t))- beta);
end

y = x/body;
plot(y)
xlabel('Time')
ylabel('Tunnel length')
```
